# A potently neutralizing and protective human antibody targeting antigenic site V on RSV and hMPV fusion glycoprotein

**DOI:** 10.1101/2024.10.31.621295

**Authors:** Alexandra A. Abu-Shmais, Luqiang Guo, Ahmed Magady Khalil, Rose J. Miller, Alexis K. Janke, Matthew J. Vukovich, Lindsay E. Bass, Yukthi P. Suresh, Scott A. Rush, Rachael M. Wolters, Nurgun Kose, Robert H. Carnahan, James E. Crowe, Rachel H. Bonami, Jarrod J. Mousa, Jason S. McLellan, Ivelin S. Georgiev

## Abstract

Human respiratory syncytial virus (RSV) and human metapneumovirus (hMPV) are frequent drivers of morbidity and mortality in susceptible populations, most often infantile, older adults, and immunocompromised. The primary target of neutralizing antibodies is the fusion (F) glycoprotein on the surface of the RSV and hMPV virion. As a result of the structural conservation between RSV and hMPV F, three antigenic regions are known to induce cross-neutralizing responses: sites III, IV, and V. Leveraging LIBRA-seq, we identify five RSV/hMPV cross-reactive human antibodies. One antibody, 5-1, potently neutralizes all tested viruses from the major subgroups of RSV and hMPV and provides protection against RSV and hMPV in a mouse challenge model. Structural analysis reveals that 5-1 utilizes an uncommon genetic signature to bind an epitope that spans sites Ø, II and V, defining a new mode of antibody cross-reactivity between RSV and hMPV F. These findings highlight the molecular and structural elements influencing RSV and hMPV cross-reactivity as well as the potential of antibody 5-1 for translational development.

## INTRODUCTION

Human respiratory syncytial virus (RSV) and human metapneumovirus (hMPV) are worldwide, endemic respiratory pathogens of the *Pneumoviridae* family^1^. Representing non-segmented negative-strand RNA viruses, RSV and hMPV induce severe and lethal bronchiolitis and pneumonia among particularly susceptible populations, most notably infantile, geriatric, and immunocompromised^2,3^, with RSV being a leading cause of lower respiratory tract infection-associated hospitalization and mortality in children under 5 years of age^4,5^. A turbulent history of disease enhancement following RSV vaccination^6^ has only recently been met with clinical success in the advancement of effective prophylactic strategies leveraging structure-based vaccine design^7–9^ and neutralizing antibodies with extended half lives^10,11^. Currently there are no approved therapeutic or prophylactic options against hMPV infection.

The major target of neutralizing antibodies in human sera against RSV and hMPV infection is the fusion (F) glycoprotein on the surface of the virion^12–15^. RSV/hMPV F is a trimeric type I transmembrane fusion protein responsible for mediating viral entry into host cells of the airway epithelium^16^. Substantial conformational changes occur in F as it transitions from the metastable prefusion form to the stable postfusion form, and understanding of these structural rearrangements has enabled engineering of prefusion-stabilized F antigens^17–21^. Stabilization of RSV and hMPV F in the prefusion state induces high neutralizing titers in experimentally inoculated animals and prefusion-stabilized RSV F serves as the backbone of the recently approved human RSV vaccines. Importantly, differential glycosylation patterns on the apex of RSV and hMPV F result in conformationally specific contributions towards the induction of neutralizing responses: RSV prefusion F epitopes are exceptionally immunogenic and invoke potently neutralizing antibodies^13,22^, whereas pre- and post-fusion hMPV F stimulate comparable neutralizing responses^20,23^. Antibody isolation and characterization efforts against RSV and hMPV have enabled extensive definition of the antigenic landscapes of RSV and hMPV F. The antigenic topology of RSV and hMPV F follows a synonymous nomenclature, with the major sites represented as site Ø through site V, as well as the more recently described site VI on RSV F^24^. Antigenic sites Ø, V and VI are preserved exclusively on the prefusion conformations of the proteins^22,25,26^, whereas sites I, II, III, and IV are exposed on the pre- and postfusion conformations.

Broadly reactive and neutralizing antibodies that recognize both RSV and hMPV have been described with varied breadth and potency of virus neutralization^22,27–34^. Due to the structural conservation between RSV and hMPV F glycoproteins, three shared epitopes on F elicit cross-reactive antibody responses, despite low sequence identity (∼35%)^35^: sites III, IV, and V. Site III is highly conserved between both viruses and a common target of cross-neutralizing antibodies encoded by *IGHV3-11/IGHV3-21: IGLV1-40*^28,32–34^, a germline gene pairing reported to be enriched in infant and adult anti-RSV antibody repertoires recognizing site III^36^. Low- and high-resolution structural analyses of site III and IV cross-reactive antibodies provide evidence that binding pose may influence cross-reactivity; however, the mode of antigenic recognition of a site V cross-neutralizing antibody remains unknown.

Leveraging LIBRA-seq (Linking B cell Receptor Sequence to Antigen Specificity by Sequencing), we identified from human PBMC samples five RSV/hMPV cross-reactive antibodies that showed high neutralization potencies against both RSV and hMPV that were comparable to virus-specific (RSV- or hMPV-only) antibodies in the literature, with one monoclonal antibody (mAb) 5-1 potently neutralizing the major subgroups of RSV and hMPV. We determined the epitope of 5-1 by single-particle cryo-EM using a prefusion-stabilized hMPV F with inter- and intra-protomer disulfide bonds and found that the binding site of 5-1 spans antigenic sites Ø, II and V on an individual protomer. Analysis of the interface identifies residues that are important for RSV and hMPV cross-neutralization. Finally, 5-1 showed robust protection in a mouse challenge model against both RSV and hMPV, therefore establishing this antibody as a prime candidate for further translational development.

## RESULTS

### Isolation of RSV/hMPV cross-reactive monoclonal antibodies by LIBRA-seq

To identify RSV/hMPV cross-reactive antibodies, we mined previously reported LIBRA-seq datasets^37,38^ that included prefusion-stabilized F glycoproteins from RSV A, RSV B, hMPV A, hMPV B, as well as control antigens. These B cells were bulk sorted from healthy donor PBMC samples, based on the expression of several markers: CD19^+^, IgG^+^, antigen^+^. After sequencing and computational filtering, we isolated a total of 27 B cells with positive signal (defined as a minimum LIBRA-seq score of 1) for at least one of the F glycoproteins belonging to both RSV and hMPV, while exhibiting low signal (defined as a LIBRA-seq score less than 1) for binding to control antigens.

### Epitope mapping and in-vitro functional properties

Five B cell receptor sequences from our analysis, corresponding to B cells with high LIBRA-seq scores of at least 1 for both RSV A/B and hMPV A/B, were produced recombinantly as IgG1 monoclonal antibodies (mAb) (Figure 1A). Four of the five antibodies are encoded by gene segments belonging to the VH3 family, with two of the four using the archetypal *IGHV3-11/3-21:IGLV1-40* of site III cross-reactive antibodies such as MPE8, 25P13, RSV199, and MxR^34^. In contrast, mAb 5-1 leveraged a pairing not yet reported, to our knowledge, among RSV/hMPV cross-reactive B cells (Figure 1B). Predicted reactivity was confirmed via enzyme-linked immunosorbent assay (ELISA) (Figure 1C). To investigate the antigenic binding sites of the cross-reactive mAbs, we tested the antibodies for competition ELISA binding against site-specific published antibodies with prefusion-stabilized RSV F and hMPV F protein antigens. Antibodies 2-6, 9-1, and 1-2 displayed consistent competition binding profiles on RSV and hMPV F proteins, mapping to sites III (2-6, 9-1) and IV (1-2). Intriguingly, mAb 5-1 strongly competed for binding to multiple sites on RSV prefusion F (sites Ø, II, III) and hMPV prefusion F (sites II, III, V). mAb 0-20 also strongly competed for binding to multiple sites on RSV prefusion F (sites Ø, II, V) and hMPV prefusion F (sites II and V) (Figure 2A). Due to the unusual competition profiles of mAbs 5-1 and 0-20, we conducted epitope binning using competition biolayer interferometry (BLI). Individually, prefusion-stabilized RSV or hMPV F protein was loaded onto sensors before saturating with mAbs 5-1 or 0-20 followed by exposure to mAbs with known antigenic epitopes. Similar to their competition ELISA binding profile, mAb 5-1 competed with site Ø, II, III, and V mAbs on RSV, and II, III, and V on hMPV, while mAb 0-20 competed with site Ø, II, and V mAbs on RSV, and site II and V mAbs on hMPV (Figure 2B).

**Figure 1:**
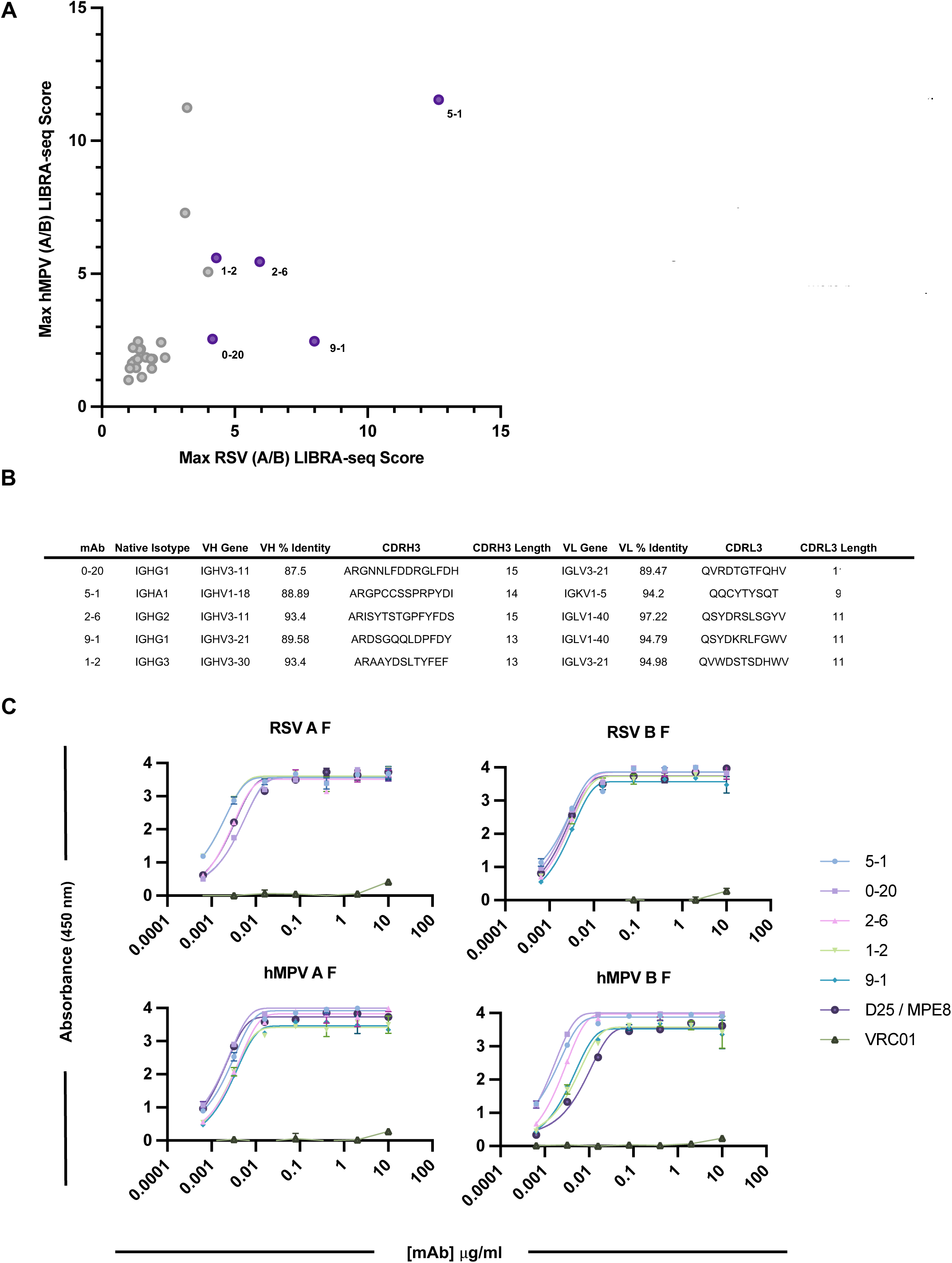
Identification and characterization of RSV/hMPV cross-reactive antibodies. A: LIBRA-seq predicted RSV and hMPV specific B cells. Each dot indicates an individual B cell. Max RSV A / RSV B LIBRA-seq score on the x-axis, max hMPV A / hMPV B LIBRA-seq score on the y-axis. Dots colored in purple were selected for further characterization. B: Sequence characteristics of RSV/hMPV cross-reactive antibodies. Percent identity is calculated at the nucleotide level and sequences and VDJ/VJ length are displayed at the amino acid level. C: ELISA binding of recombinantly produced antibodies against RSV and hMPV prefusion F trimer, calculated as absorbance at 450 nm. Experiments were performed in technical and biological duplicate.

**Figure 2:**
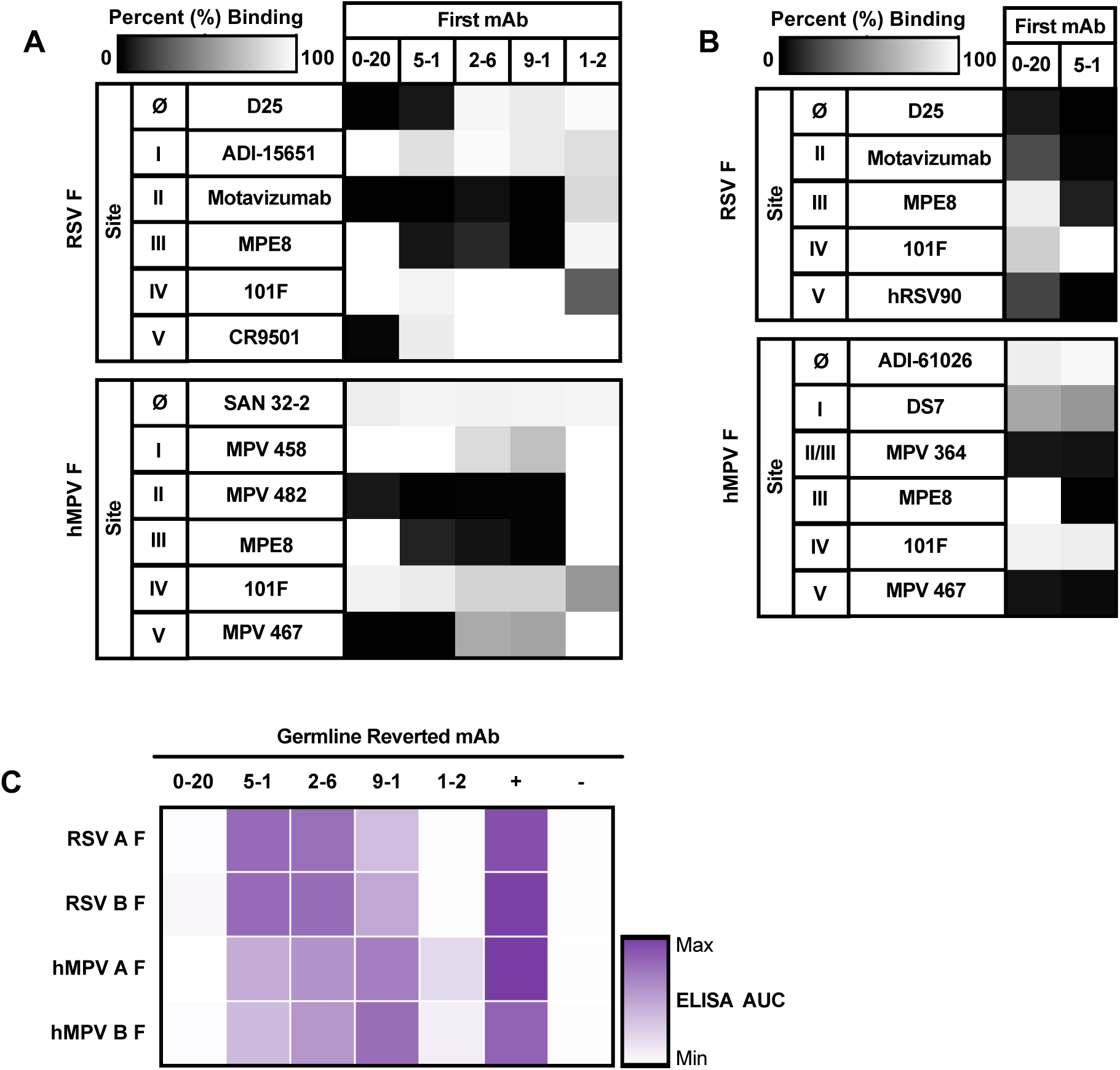
Binding characteristics of RSV/hMPV cross-reactive mAbs. A: Antibody-antibody competition binding to RSV and hMPV prefusion F trimer against control site specific antibodies. Percentage of binding of biotinylated antibody is shown as a heatmap from 0% (black) to 100% (white). Non-biotinylated competitor antibodies were coated first, and then biotinylated control mAbs were added to detect competition. Competition is calculated as the signal obtained for binding of the biotin-labelled reference antibody in the presence of the unlabeled antibody, expressed as a percentage of the binding of the reference antibody alone. B: Epitope binning via BLI for binding of mAbs 20 and 5-1 to RSV and hMPV prefusion F trimer. Data indicate the percent binding of the second antibody in the presence of the first antibody, as compared to the second antibody alone. Percentage of binding is shown as a heatmap from 0% (black) to 100% (white). C: ELISA binding of germline reverted, recombinantly produced antibodies against RSV A and B and hMPV A and B prefusion F trimer, calculated as absorbance at 450 nm. ELISA area under the curve (AUC) shown as a heatmap from minimum (white) to maximum binding (purple).

To investigate whether cross-reactivity emerged as a result of somatic hypermutation, we reverted each candidate mAb to its germline sequence and tested binding to recombinant F antigens. While mAbs 9-1 and 2-6 both target site III, germline-reverted mAb 2-6 preferred binding to RSV F while germline-reverted mAb 9-1 preferred binding to hMPV F (Figure 2C). Binding to both RSV and hMPV F was abrogated for the germline-reverted mAb 0-20, while mAb 5-1 and mAb 1-2 displayed preferential binding to RSV F and hMPV F, respectively (Figure 2C).

Antibody-virus neutralization potency was determined by plaque reduction neutralization test (PRNT) using live virus to inoculate cells. All candidate mAbs exhibited neutralization against at least one of the tested viruses representing the major antigenic groups of RSV and hMPV. Notably, while mAb 5-1 demonstrated higher neutralization potencies against hMPV compared to RSV viruses, this antibody exhibited strong neutralization against all viruses tested (IC50 0.0029–0.0280 µg/mL) (Figure 3A-B). To assess autoreactivity, binding to permeabilized HEp-2 cells was performed. At 1 µg/mL and 10 µg/mL, none of the antibodies displayed binding to HEp-2 cells (Supplementary Figure 1).

**Figure 3:**
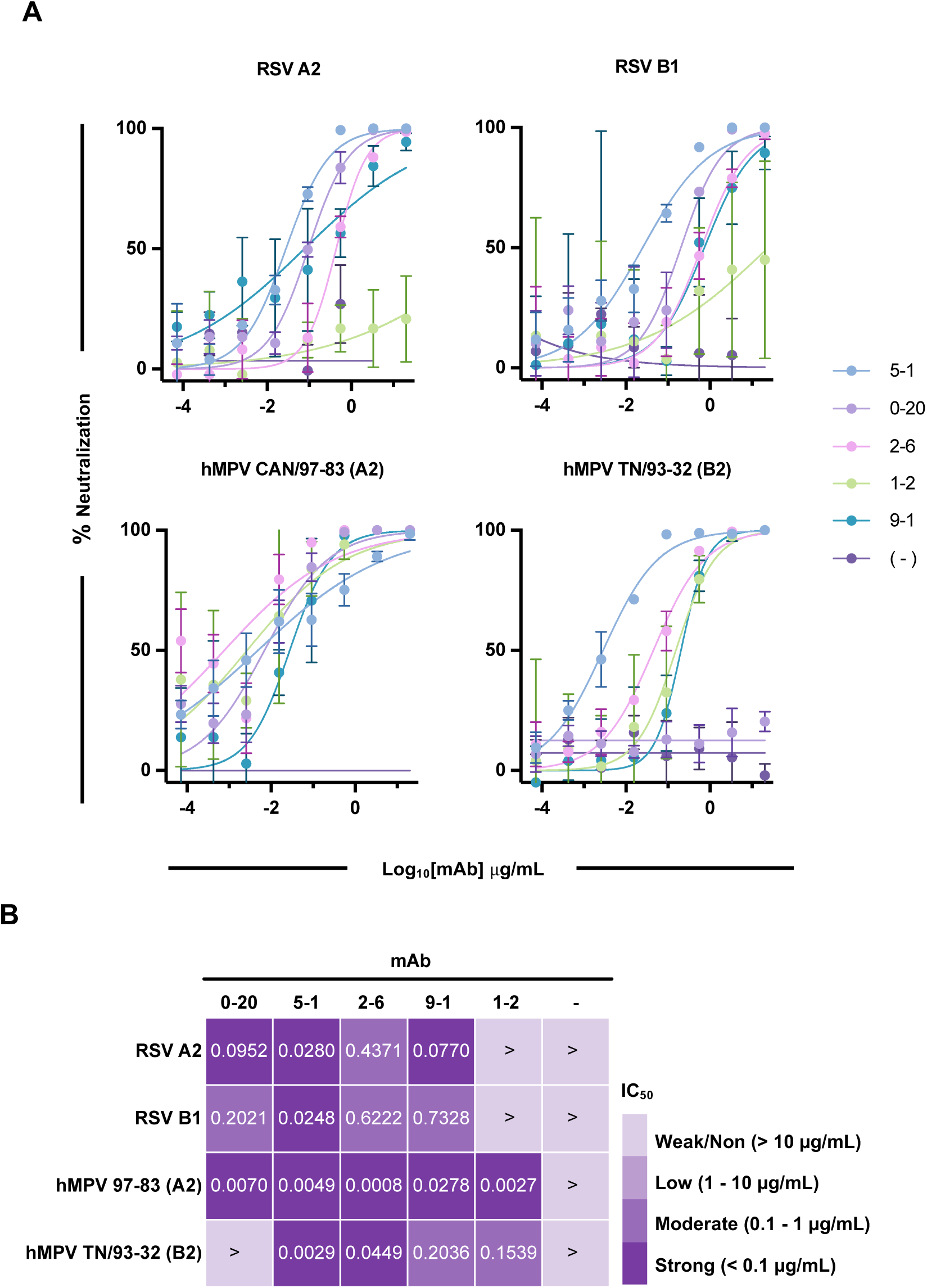
Neutralization potency of RSV/hMPV cross-reactive mAbs. A: Antibody neutralization against RSV A2, RSV B1, hMPV A2, and hMPV B2 via PRNT. B: IC50 values, expressed as a heatmap with strong neutralization (<0.1 µg/mL) shown in purple and weak/non neutralizing (>10 µg/mL) shown in light purple. Calculated by non-linear regression analysis by GraphPad Prism software. Neutralization assays were performed in technical triplicate; data are represented as mean ± SD.

### Structure of mAb 5-1 complexed with hMPV F

As mAb 5-1 was the most potently neutralizing antibody and displayed a unique competition profile that was not resolved by competition biolayer interferometry, we investigated the epitope of mAb 5-1 using negative stain electron microscopy (EM) and cryo-electron microscopy (cryoEM). Efforts with a prefusion RSV F protein (DS-Cav1) and 5-1 antigen-binding fragment (Fab) were unsuccessful, as most of the trimers were observed in a splayed-open state (Supplementary Figure 2). Therefore, we used a prefusion-stabilized hMPV F construct (hMPV F-DS-CavEs2-IPDS), which contains intra- and inter-protomer disulfide bonds to lock hMPV F in a closed prefusion trimer conformation^39^

Cryo-EM analysis of hMPV F and 5-1 Fab revealed a heterogeneous mixture of complexes composed of three Fabs per trimer, with the majority of the particles displaying flexibility at the membrane-proximal base of the F protein (Supplementary Figure 3). However, a subset of particles retained after 2D classification were identified with a well-ordered base (∼23%), and further processing yielded a 3D reconstruction with a global resolution of 4.3 Å (Supplementary Figure 4B,C). The cryo-EM map agrees very well with a model of the complex produced with AlphaFold3^40^, and only light refinement was required to obtain an excellent map-to-model fit.

The structure reveals that the 5-1 epitope is contained within the F1 subunit of a single protomer and primarily spans antigenic sites II and V, with some additional interactions with site Ø (Figure 4A,B). The 5-1 heavy and light chains bury 597 Å^2^ and 303 Å^2^ of surface area, respectively, with the complementarity-determining region (CDR) 1 and 2 of the light chain contributing to the interaction with site Ø and the top half of site V. The light chain primarily interacts with residues on α4 through an electrostatic interaction network formed by Asp31CDRL1 and Arg50CDRL2 with RSV F residues Lys171 and Asp167, and with residues on the loop preceding β3 through the electrostatic interaction of Glu55CDRL2 with Lys143 (Figure 4C). The heavy chain packs its CDR loops against the cleft between β3 and α6, with Tyr53CDRH2 inserted into the cleft. Interestingly, the 5-1 CDRL3 only interacts with residues on the CDRH2 and CDRH3 loops rather than with hMPV F, which may be important for stabilizing the heavy chain interactions (Figure 4C). In addition, there appear to be interactions between light chain framework residues and the N-linked glycans attached to Asn172 on hMPV F, despite the low resolution and partially modeled glycan chains (Supplementary Figure 4A-D).

**Figure 4:**
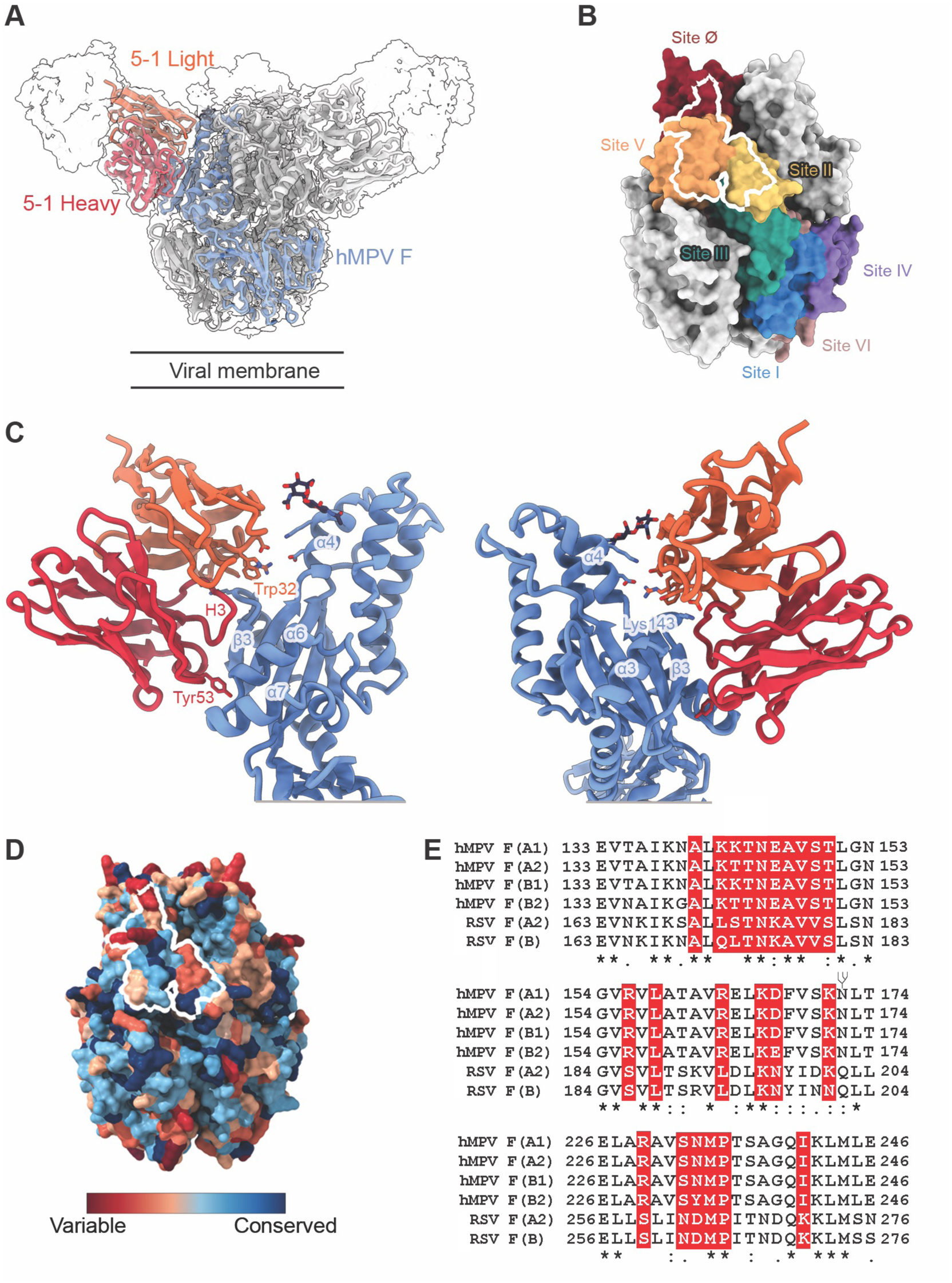
5-1 Fab binds to the prefusion hMPV F at site II, V and the glycan at Asn172. A: Front view and side view of the fit of hMPV F complex into a DeepEMhanced EM map at the contour level of 0.432. The global DeepEMhanced EM map was show as a white transparent map with a single hMPV F protomer and Fab variable domain colored (hMPV F, blue; heavy chain variable domain, red; light chain variable domain, orange). B: Overlay of the 5-1 epitope onto the defined antigenic sites of hMPV F revealing that 5-1 primarily interacts with residues in site II and V, with additional contacts within site Ø. C: Atomic model of 5-1 and hMPV F interface with key residues highlighted as sticks. 5-1 and one hMPV F protomer are shown as cartoons. Oxygen atoms are colored red and Nitrogen atoms are colored blue. Partially modeled Asn-172 glycan is shown as deep color sticks. D: Sequence conservation of the 5-1 epitope between hMPV F and RSV F with the epitope of 5-1 delineated in white. E: Sequence alignment of the 5-1 epitope with four representative hMPV F sequences from A1, A2, B1, B2 subgroup and two representative RSV sequences from A2 and B subgroup. The conservation of each residue is described underneath and the 5-1 interacting residues are highlighted in red. The glycosylation site at Asn-172 is shown as a branch.

The structural model obtained from cryo-EM analysis agrees well with the ELISA and BLI competition binding data. Superposition of the cryo-EM structure with previously determined structures of the antibodies used in the competition assays predicts that 5-1 would sterically clash with D25, motavizumab, MPE8, hRSV90, ADI-61026 and MPV467 (Supplementary Figure 5). Further comparison to known hMPV and RSV F antibody complexes revealed that hRSV90 binds to a similar epitope on RSV F, except with an inverted arrangement of the heavy and light chains (Supplementary 6). However, hRSV90 is specific for RSV and does not bind or neutralize hMPV.

The 5-1 epitope contains some amino acids that are not well conserved among RSV and hMPV F proteins, yet the antibody binding mode can accommodate these differences (Figure 4D,E). The substitutions will likely impact the affinity of 5-1 to different extents, but they do not introduce clashes that would prevent antibody binding. The region including the β3 strand is generally well conserved (hMPV F residues 142–150), as is the cleft between β3 and α6, into which Tyr53CDRH2 inserts. Thus, the structure and AlphaFold3 models of 5-1 bound to hMPV F and RSV F provide a structural basis for how 5-1 can bind an epitope at the F apex that is thought to be under immune pressure and less conserved than other regions.

### In-vivo protection against viral infection

Next, we investigated the protective efficacy of mAb 5-1 in both an RSV and hMPV infection model in BALB/c mice. Fourteen-week-old female mice were mock treated with PBS, an isotype control human mAb VRC01, or different doses of mAb 5-1 six hours prior to intranasal RSV or hMPV challenge (Figure 5A,B). Lung viral titers of mice were determined by plaque assay on day 6 post infection to assess mAb 5-1 prophylaxis against infection. At the highest mAb 5-1 dose of 10 mg/kg, viral lung titers were below the detection limits for both RSV and hMPV for all animals (Fig. 5B). Even at the 10-fold lower dose of 1 mg/kg, 2/5 animals (40%) showed no detectable viral titers in the lung for both RSV and hMPV and were overall significantly lower than those observed in the control groups. Animals receiving the lowest dose of 0.1 mg/kg of mAb 5-1 showed significantly reduced lung viral titers for RSV and a 3.33-fold (though not statistically significant) reduction for hMPV. Together, these results showcase the *in vivo* protective ability of mAb 5-1 against RSV and hMPV challenge.

**Figure 5:**
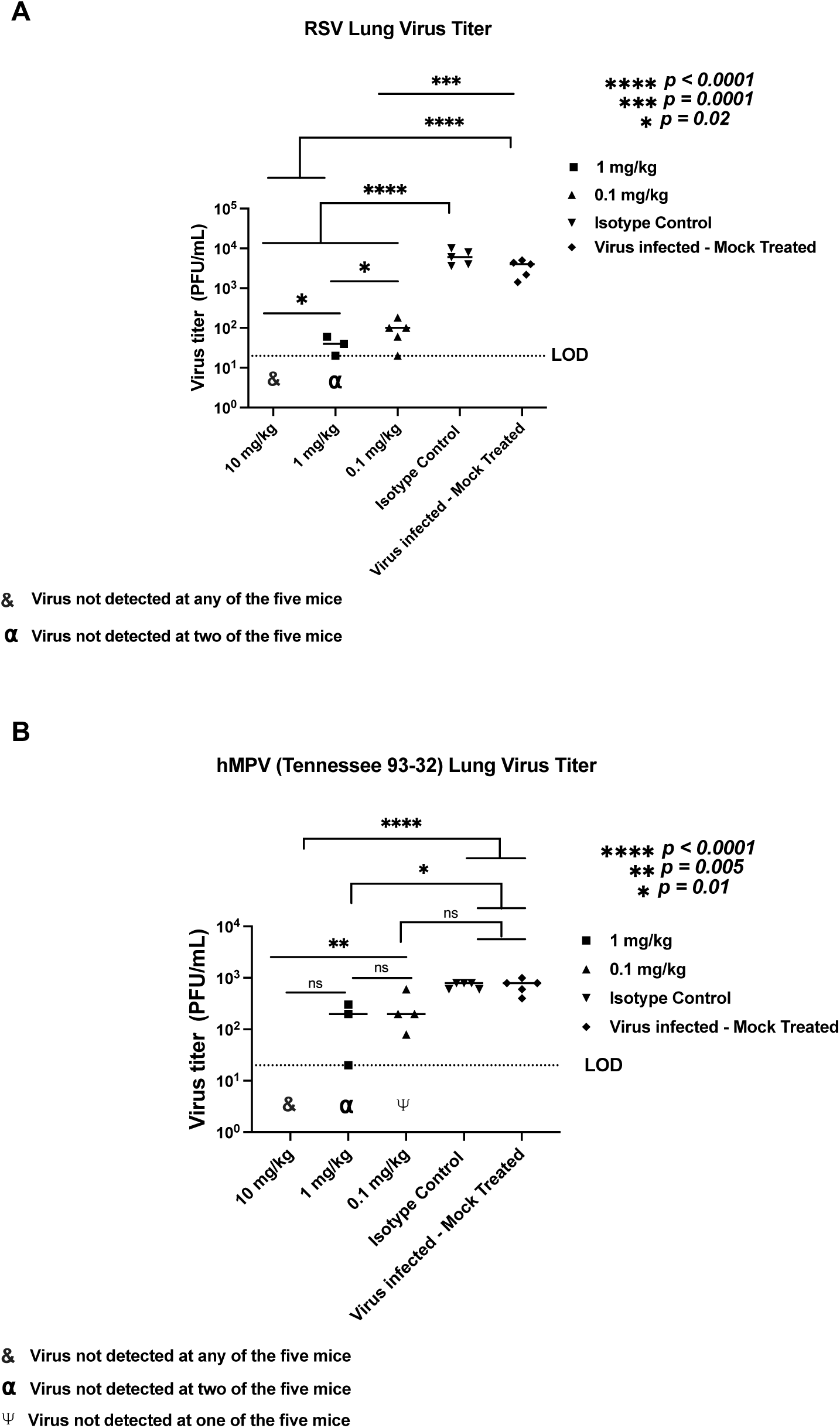
5-1 Prophylaxis of 5-1 against RSV and hMPV challenge. Protective efficacy of 5-1 against A) RSV and B) hMPV replication in *vivo*. BALB/c mice were treated intraperitoneally with 10 mg/kg, 1 mg/kg, and 0.1 mg/kg of mAb 5-1 6h prior to intranasal RSV and hMPV infection. Viral titers in the lung homogenates of BALB/c mice in each treatment group (n = 5 mice per group, 5 females) were determined by plaque assay. n.s., not significant, Limit of detection (LOD) is indicated with a dashed line.

## DISCUSSION

Respiratory illness associated with infection by either RSV and/or hMPV remains a public health threat, with the potential for severe disease in neonatal, geriatric, and immunocompromised patients such as those undergoing hematopoietic stem cell transplant and patients suffering from pulmonary co-morbidities. While strategies to prevent severe infection induced by RSV have advanced in the last year, there are currently no approved treatments for infection by hMPV. We and others have isolated RSV and hMPV cross-neutralizing antibodies that present an interesting alternative to mono-valent therapies, providing a protective regimen for the prevention or amelioration of disease caused by either mono- or co-infection of RSV and hMPV.

We discovered five antibodies targeting three previously reported epitopes on the F protein known to elicit cross-reactive humoral responses. Consistent with the enrichment of site III-directed antibodies encoded by *IGHV3-11/3-21*:*IGLV1-40*, mAbs 9-1 and 2-6 display competition profiles indicative of binding at antigenic site III. Interestingly, germline-reverted mAbs 9-1 and 2-6 favored binding to F from different viruses, despite targeting the same site. Loss of antigenic binding of mAb 0-20 to both RSV and hMPV F in the germline state suggests cross-reactivity can be achieved through multiple antibody evolution pathways, i.e., through subsequent activation of either RSV or hMPV-specific B cells.

All five RSV/hMPV antibodies displayed *in vitro* neutralizing activity against infection by at least one representative virus of each genotype, albeit some mAbs displayed preferential neutralization against RSV or hMPV alone. mAb 5-1 displayed potent neutralization against all viruses tested, reaching neutralization potencies of better than 10 ng/mL IC50 against hMPV 97-83 and hMPV TN/93-32. A significant proportion of hMPV field strains contain amino acid substitution D280N^41,42^, which may impede binding of *IGHV3-11/3-21:IGLV1-40* site III cross-reactive antibodies such as MPE8, 25P13, and RSV199. Our structural analysis predicts this mutation would be well tolerated, as D280 falls outside of the epitope of 5-1, which is predominantly within antigenic site V and antigenic site II, with additional contacts with site Ø. The structure agrees well with the ELISA and BLI competition assay data, with the exception for antibody DS7. The modeling indicates that DS7 is not predicted to clash with 5-1, however some competition was observed (Figure 2B). This may be influenced by the ability of DS7 to bind a conformation of the hMPV F protomer that contains elements of both the prefusion and postfusion conformation ^43^. The apex of hMPV F is shielded by glycans on Asn57 and Asn172 (Supplementary Figure 7), reducing antigenic exposure and dampening the immune response, relative to that of RSV, against site V and site Ø ^23,44–46^. However, despite this immune evasion technique, the human immune system has proven its ability to circumvent this obstacle through penetration of the glycan shield, as demonstrated with antibody ADI-61026^44^, where ADI-61026 positions itself into a pocket between two glycans and directly interacts with Asn57-glycan. Glycan-shield-penetrating antibodies have also been reported that bind to HIV-1 Env^47,48^, and hepatitis C E2 ^49^. Here, we demonstrate that 5-1 is also able to breach the glycan shield at the apex of hMPV F (Supplementary Figure 7).

As mAb 5-1 predominantly targets antigenic site V and provides protection against hMPV and RSV infection, we systematically compared the 5-1 binding pose and epitope with other site V antibodies, where structural information was available. We observed that antibodies bind to site V with varied modes of binding and thus contact differential residues in their respective epitopes, as observed with site III^28,33,34^ and IV binders^29^. While many of the antibodies discussed here engage site V contact residues that are conserved between hMPV and RSV, the majority of these antibodies retain specificity for RSV or hMPV alone, likely as a result of the structural difference between RSV and hMPV trimers (Supplementary Figure 6).

Structural and repertoire analyses, in the context of antibodies elicited as a result of natural infection by RSV and hMPV^22,39,45,50,51^, have revealed the propensity of site V towards the induction of potently neutralizing humoral responses. Within the trimeric prefusion F protein, the fusion peptide is buried inside a hydrophobic cavity occluded by the site V epitope. As demonstrated with a previously reported antibody targeting site V on hMPV F^39^, one potential explanation for the potency of mAb 5-1, as compared to the other mAbs in our set, is that binding of mAb 5-1 prevents extension of the fusion peptide from the F protein, thereby disrupting the conformational changes necessary for productive infection.

Currently, no FDA-approved prophylaxis or therapeutics against hMPV F are available, despite substantial efforts^52,53^. Recent progress, including structure-based RSV vaccines and antibody prophylaxis have been made, yet an antibody that potently neutralizes RSV with a unique antigenic footprint may offer additional benefits when considering the potential for virus evolution. Furthermore, an antibody that provides cross protection against both RSV and hMPV infection can be utilized to provide long-lasting protection against infection from either of these viruses in at-risk populations, providing important logistical advantages over developing multiple virus-specific mAbs. mAb 5-1 therefore presents an attractive target for further translational development.

## MATERIALS AND METHODS

### Data mining

LIBRA-seq datasets generated from 2020-2023 that included prefusion RSV A F, RSV B F, hMPV A F, and hMPV B F in the antigen screening library were mined for B cells displaying a minimum LIBRA-seq score of one for at least one of the F antigens, while also displaying a score below one for a control antigen, in this case, recombinant HIV-1 envelope protein. LIBRA-seq experiments were performed on peripheral blood mononuclear cells (PBMCs) samples obtained from otherwise healthy adult individuals. The established LIBRA-seq pipeline was used for score generation^54^.

### Antibody expression and purification

For each antibody, variable genes were synthesized as cDNA and were inserted into bi-cistronic plasmids encoding for the constant regions of the heavy chain and either the kappa or lambda light chain (Twist BioScience). Antibodies were transiently expressed with Expifectamine transfection reagent (Thermo Fisher Scientific) in Expi293F cells in FreeStyle F17 expression media (Thermo Fisher) (0.1% Pluronic Acid F-68 and 20% 4 mM L-glutamine). Cells were cultured for 5 days at 8% CO2 saturation and 37°C with shaking. Five days post transfection, cells were collected and centrifuged at a minimum of 6000 rpm for 20 minutes. Supernatant was filtered with Nalgene Rapid Flow Disposable Filter Units with PES membrane (0.45 or 0.22 μm) and purified over protein A equilibrated with PBS. Antibodies were eluted with 100 mM glycine HCl at pH 2.7 directly into a 1:10 volume of 1 M Tris-HCl pH 8 and then exchanged into PBS for storage at 4°C.

### Enzyme linked immunosorbent assay (ELISA)

Recombinant antigen was plated at 2 ug/mL overnight at 4°C. The next day, plates were washed three times with PBS supplemented with 0.05% Tween20 (PBS-T) and coated with 1% bovine serum albumin (BSA) in PBS-T. Plates were incubated for one hour at room temperature and then washed three times with PBS-T. Primary antibodies were diluted in 1% BSA in PBS-T, starting at 10 μg/mL with a serial 1:5 dilution, plated, and then incubated at room temperature for one hour before washing three times in PBS-T. The secondary antibody, goat anti-human IgG conjugated to peroxidase, was added at 1:10,000 dilution in 1% BSA in PBS-T to the plates, which were incubated for one hour at room temperature. Plates were washed three times with PBS-T and then developed by adding TMB substrate to each well. The plates were incubated at room temperature for five minutes, and then 1 N sulfuric acid was added to stop the reaction. Plates were read at 450 nm. ELISAs were performed in technical and biological duplicate.

### Competitive binding of mAbs with site-specific antibodies in the literature

Wells of 384-well microtiter plates were coated with 25ul of 2 μg/mL purified F antigenic protein at 4°C overnight. Plates were blocked with 50 μl of 1% BSA in PBS-T for 1 h before washing three times with PBS-T. Primary antibodies at 10 μg/mL were added to wells (20 μL per well) in duplicate and incubated for 1 h at room temperature. A biotinylated preparation of recombinantly produced site-specific monoclonal antibodies were added to wells of each primary antibody at a concentration of 10μg/mL in a volume of 5 μL per well, without washing of unlabeled antibody, and then incubated for 1 h at room temperature. Plates were washed three times with PBS-T and bound antibodies were detected using horseradish peroxidase (HRP) -conjugated anti-biotin 1:1000 (ThermoFischer Scientific) and a TMB substrate. The signal obtained for binding of the biotin-labelled reference antibody in the presence of the unlabeled tested antibody was expressed as a percentage of the binding of the reference antibody alone after subtracting the background signal. Tested mAbs were considered competing if their presence reduced the reference antibody binding to less than 40% of its maximal binding and non-competing if the signal was greater than 71%. A level of 41 to 70% was considered intermediate competition.

### Germline Reversion of BCRs

Nucleotide sequences for the heavy and light chains of the described antibodies were annotated using IMGT V-Quest. Mutations occurring outside of the CDR3 region were reverted to the residues present in the V and J genes and alleles that most closely aligned to the mature sequence.

### Cell culture and virus CPE determination

LLC-MK2 cells were obtained from ATCC (CCL-7) and grown in growth media (Opti-MEM with 2% FBS) at 37°C, 5% CO2. Propagated virus was grown in viral growth media (Opti-MEM with 5 µg/mL trypsin-EDTA and 1% antibiotic-antimycotic) in LLC-MK2 cells at a multiplicity of infection (MOI) of 0.01 for 3-5 days at 37°C, 5% CO2 until CPE was observed. Virus was harvested using the freeze-thaw method into 25% sucrose solution and stored at -80°C until use.

### Plaque reduction neutralization test with MPV (CAN/97-83 and TN/93-32) or RSV (A2 and B) virus

24 hours prior to viral infection, LLC-MK2 (for hMPV) or HEp-2 (for RSV) cells were plated in growth media at 5 × 10^4^ cells per well in 24 well plates and incubated at 37°C, 5% CO2. The day of viral infection, mAbs were serially diluted in Opti-MEM with a starting concentration of 40 µg/mL. hMPV (CAN/97-83 and TN/93-32) or RSV (A2 and B) virus was diluted in Opti-MEM to a final concentration of 2400 plaque forming units (pfu)/mL and added to the mAb mixtures at a 1:1 volume ratio. The mAb/virus mixture incubated for 1 hour at room temperature. Prior to adding the mAb/virus mixture to cells, confluent cells in 24 well plates were washed gently three times with PBS. mAb/virus mixture was added to each well (50 µL per well) and the plates rocked at 37°C, 5% CO2 for 1 hour. Warm overlay (0.75% methylcellulose in Opti-MEM, 5 µg/mL trypsin-EDTA and 1% antibiotic-antimycotic) was added to each well and the plates incubated for 4 days at 37°C, 5% CO2. Following incubation, the cells were fixed with 10% neutral buffered formalin, washed with water three times, then blocked with milk blocking buffer (2% milk powder, 2% goat serum in PBS-T). Plates were washed three times with water and immunostained with human mAbs MPV364 (for hMPV) or 101F (for RSV) diluted to 5 µg/mL in milk blocking solution for 1 hour at room temperature. Plates were washed three times with water before adding the secondary antibody, goat anti-human IgG Fc conjugated to horse radish peroxidase, at a dilution of 1:2000 in milk blocking solution and incubated for 1 hour at room temperature. Plates were washed three times with water and developed with TrueBlue substrate by rocking for 10 minutes. After plaques were visibly stained by the substrate, the plates were washed once with water to stop the developing reaction. Immunostained plaques were counted and graphed on GraphPad Prism9.

### RSV and hMPV mouse challenge model

BALB/c mice (14 weeks old; The Jackson Laboratory) were intranasally infected with RSV A2 (2.0E+6 PFU/mouse) or hMPV TN/93-32 (3.0E+5 PFU/mouse) and euthanized 6 d postinfection. Monoclonal antibody 5-1 was administered intraperitoneally at 10, 1.0, or 0.1 mg/kg. Control mice were intraperitoneally injected with PBS or VRC01 (isotype control) at 10 mg/kg. All injections occurred 6 h prior to infection. Lung homogenates were used for viral titration by plaque assay as described above.

### HEp-2 cell immunofluorescence assay to detect mAb autoreactivity

HEp-2 cell coated slides (BION ENTERPRISES LTD ANA (Hep-2) Test System, ANK-120) were incubated with purified antibodies at 10 and 1 ug/ml or control sera in a moist chamber at room temperature for 30 min. Controls provided with the kit included anti-nuclear antibody (ANA)^+^ and (ANA)^-^ human sera. Slides were washed twice with PBS for 5 min. Cells were stained with FITC-goat anti-human Ig per the manufacturer’s instructions and incubated in a moist chamber at room temperature for 30 min. Slides were washed twice with PBS for 5 min, mounted with DAPI mounting medium (Southern Biotech 0100-20) and visualized by fluorescence microscopy (Olympus BX60 epifluorescence microscope coupled with a CCD camera and MagnaFire software Optronics International) at 40x magnification. Image brightness and contrast were optimized using Adobe Photoshop.

### Recombinant protein production for negative stain and cryo-EM

Prefusion RSV-F strain A2 (DS-Cav-1)^17,55^ was used for negative stain-EM. Prefusion hMPV-F construct DS-CavEs2-IPDS protein was used for cryo-EM structural studies as previously reported^39^. In brief, plasmids encoding antigens were transfected into FreeStyle 293F cells (ThemoFisher) by PEI. Kifunensine and Pluronic F-68 (Gibco) were introduced 3 h post transfection. Six days later, the cell supernatant was filtered, and buffer exchanged into PBS by tangential flow filtration. Then, Step-TactinXT 4 Flow resin (IBA) was used to purify the protein from the filtered supernatant following the manufacturer’s instruction. The purified protein was then concentrated using a 10 kDa molecular weight cutoff Amicon Ultra-15 centrifugal filter unit (Millipore) and subject to a Superose 6 increase 10/300 column (Cytivia) in PBS running buffer (hMPV-F DS-CavEs2-IPDS) or 2 mM Tris pH 8.0, 200 mM NaCl, and 0.02% NaN3 (RSV A2 DS-Cav-1) for preparative size-exclusion chromatography. Peaks corresponding to trimeric species were identified based on elution volume and SDS-PAGE of elution fractions. Fractions containing pure fusion protein were pooled.

### Negative stain-EM

For screening and imaging of negatively stained 5-1 Fab in complex with RSV-F A2 DS-Cav-1, sample was diluted to 100mg/mL with buffer containing 10 mM NaCl, 20 mM HEPES buffer, pH 7.4, and 5% glycerol and applied to glow-discharged grid with continuous carbon film on 400 square mesh copper EM grids (Electron Microscopy Sciences). The grids were stained with 2% uranyl formate (UF). Grids were examined on a 100 kV Morgagni microscope with a 1k x 1k AMT CCD camera.

### Cryo-EM sample preparation and data collection

The purified hMPV-F-DS-CavEs2-IPDS was combined with 5-1 Fab in PBS buffer with a final concentration of 4.8 μM and 21.6 μM and incubated on ice for 3 min. Then, the 3 μl mixture was applied to a UltrAuFoil R1.2/1.3 300 mesh grid (Electron Microscopy Sciences) that had been glow-discharged with a PELCO easiGlow glow discharge cleaning system for 1 min. Grids were plunge-frozen using a Vitrobot Mark IV (ThermoFisher Scientific) at 4 °C, 100% humidity. Blot settings were 4s of blotting with force 2. Movies (3,538) were collected from a single grid on a 200 kV Glacios microscope (ThermoFisher Scientific) equipped with a Falcon 4 direct electron detector (ThermoFisher Scientific). Data were collected at a 50-degree tilt and at a magnification of 150,000x, where the calibrated pixel size is 0.94 Å/pix and the total exposure is 48.6 e^-^/Å^2^.

### Cryo-EM data processing

Movies were imported into cryoSPARC v4.4.0^56^ for gain correction, motion correction, patch CTF estimation, micrograph curation, particle picking, and particle extraction with a 2X Fourier crop. After two rounds of particle curation through 2D class averaging, the generated 2D class averages were used as templates to perform another round of template-based particle picking. Then, the particles were curated by 2D class averaging and curated particles were subject to ab initio reconstruction, heterogeneous refinement, and homogeneous refinement with C3 symmetry applied. Due to the presence of flexibility at the bottom region of the homogeneous-refined EM map, a 3D variability analysis job was performed with a focused mask to explore alternative conformations. After 3D variability analysis, a 3D classification job with a focused mask on the hMPV F base region was executed to generate EM maps of different conformations, followed by heterogeneous refinement. As particles were processed with Fourier cropping in the procedure described above, we re-extracted the particles with raw pixel size, removed the duplicate particles and reconstructed one EM map with homogeneous refinement and reference-based motion correction. Finally, the map from the last round of homogeneous refinement was sharpened using DeepEMhancer^57^. For model building, an initial model was generated by AlphaFold3 server^58^. As the predicted model aligned well with our 3D EM map, the following iterative refinements were performed using this model in Coot^59,60^, PHENIX ^61^ and ISOLDE^62^. The adjacent cystines in 5-1 Fab CDRH3 loop were modeled as a disulfide bond in the AlphaFold3 predicted model and were left unchanged during refinement. At the last round of refinement, glycans were built into the model, refined and validated using Coot and Privateer software^63^. The EM processing workflow is shown as Supplementary Figure 3 and EM validation results are shown in Supplementary Figure 4. Refinement statistics are shown in Supplementary Table 1.

### Sequence conservation analysis and alignment

The glycoprotein sequence of hMPV F protein from strain NL/1/100 (A1 sub lineage, NCBI accession: YP_009513268.1) was uploaded into the HMMER web server^64^ to search for homologous sequences against UniProtKB database with phmmer programs and default parameters. The searching results were then manually filtered based on species, similarity, coverage and hit position. To avoid potential bias, 250 sequences for both hMPV F and RSV F were extracted from the search results and aligned with Clustal Omega^65^. The output was imported into ChimeraX^66^ to generate a sequence conservation map. For direct alignment of four representative hMPV F and two RSV F protein sequences, hMPV F from A1 (NL/1/00 strain, NCBI accession: NC_039199.1), A2 (NL/17/00 strain, NCBI accession: AAQ90144.1), B1 (NL/1/99 strain, NCBI accession: AAQ90145.1), B2 (NL/1/94 strain, AAQ90146.1) and RSV F from A2 (NCBI accession: ACO83301.1) and B (NCBI accession: WKU63582.1) sequences were pooled and aligned with Clustal Omega.

### Data Availability

Datasets from which individual antibody sequences were pulled can be found in ^37,38^. The EM map and coordinates for the hMPV F and 5-1 Fab complex have been deposited into the Electron Microscopy Data Bank (EMDB-45412) and the Protein Data Bank (9CB1; DOI: https://doi.org/10.2210/pdb9CB1/pdb). All data are included in the article and/or supporting information.

## ACKNOWLEDGMENTS

We thank all members of the Georgiev laboratory for their support and feedback. We thank David Flaherty, Olivia Murfield, Emma McLaughlin, and Brittany Matlock from the VUMC Flow Cytometry Shared Resource for their help with cell sorting. The VUMC Flow Cytometry Shared Resource is supported by the Vanderbilt Ingram Cancer Center (P30 CA68485) and the Vanderbilt Digestive Disease Research Center (DK058404). We thank Angela Jones, Jamie Roberson, and Latha Raju with the Vanderbilt Technologies for Advanced Genomics Core (VANTAGE) for providing technical assistance with library production and sequencing. VANTAGE is supported in part by CTSA (5UL1 RR024975-03), the Vanderbilt-Ingram Cancer Center (P30 CA68485), the Vanderbilt Vision Center (P30 EY08126), and NIH/NCRR (G20 RR030956). For work described in this manuscript, I.S.G., A.A.A., A.K.J, and M.J.V. were supported in part, by the G. Harold and Leila Y. Mathers Charitable Foundation (MF-2107-01851) and NIH R01AI175245 (to I.S.G.). L.G., S.A.R and J.S.M were supported in part by Welch Foundation (F-0003-19620604). A.A.A. and R.M.W. were supported in part by NIH grant T32 (5T32AI112541-07). R.M.W. was supported by 1K01OD036063-01. L.E.B. was supported by NIH T32 AR059039 and F31 DK141224. R.H.B. was supported by NIH R01 DK131070. The funders had no role in the conceptualization or execution of any studies or drafting of the manuscript.

## AUTHOR CONTRIBUTIONS

Conceptualization and Methodology: A.A.A. and I.S.G.; Investigation: A.A.A., L.G., A.K., R.J.M., A.K.J., M.J.V., L.E.B., S.A.R., Y.P.S., R.M.W, N.K.; Writing – Original Draft: A.A.A. and I.S.G.; Writing – Review & Editing: All authors; Funding Acquisition: A.A.A., J.S.M., and I.S.G. Resources: J.S.M, J.J.M., R.H.B., R.H.C., J.E.C., I.S.G; Supervision: A.A.A., J.S.M., J.J.M., R.H.B., R.H.C., J.E.C., and I.S.G.

## DECELERATION OF INTERESTS

A.A.A. and I.S.G. are listed as inventors on patents filed describing the antibodies discovered here. I.S.G. is listed as an inventor on patent applications for the LIBRA-seq technology. I.S.G. is a co-founder of AbSeek Bio. I.S.G. has served as a consultant for Sanofi. The Georgiev laboratory at VUMC has received unrelated funding from Merck and Takeda Pharmaceuticals. J.E.C. has served as a consultant for Luna Labs USA, Merck Sharp & Dohme Corporation, Emergent Biosolutions, a former member of the Scientific Advisory Boards of Gigagen (Grifols), of Meissa Vaccines, and BTG International, is founder of IDBiologics and receives royalties from UpToDate. The laboratory of J.E.C. received unrelated sponsored research agreements from AstraZeneca, Takeda Vaccines, and IDBiologics during the conduct of the study.

---

**Supplementary Figure 1:**
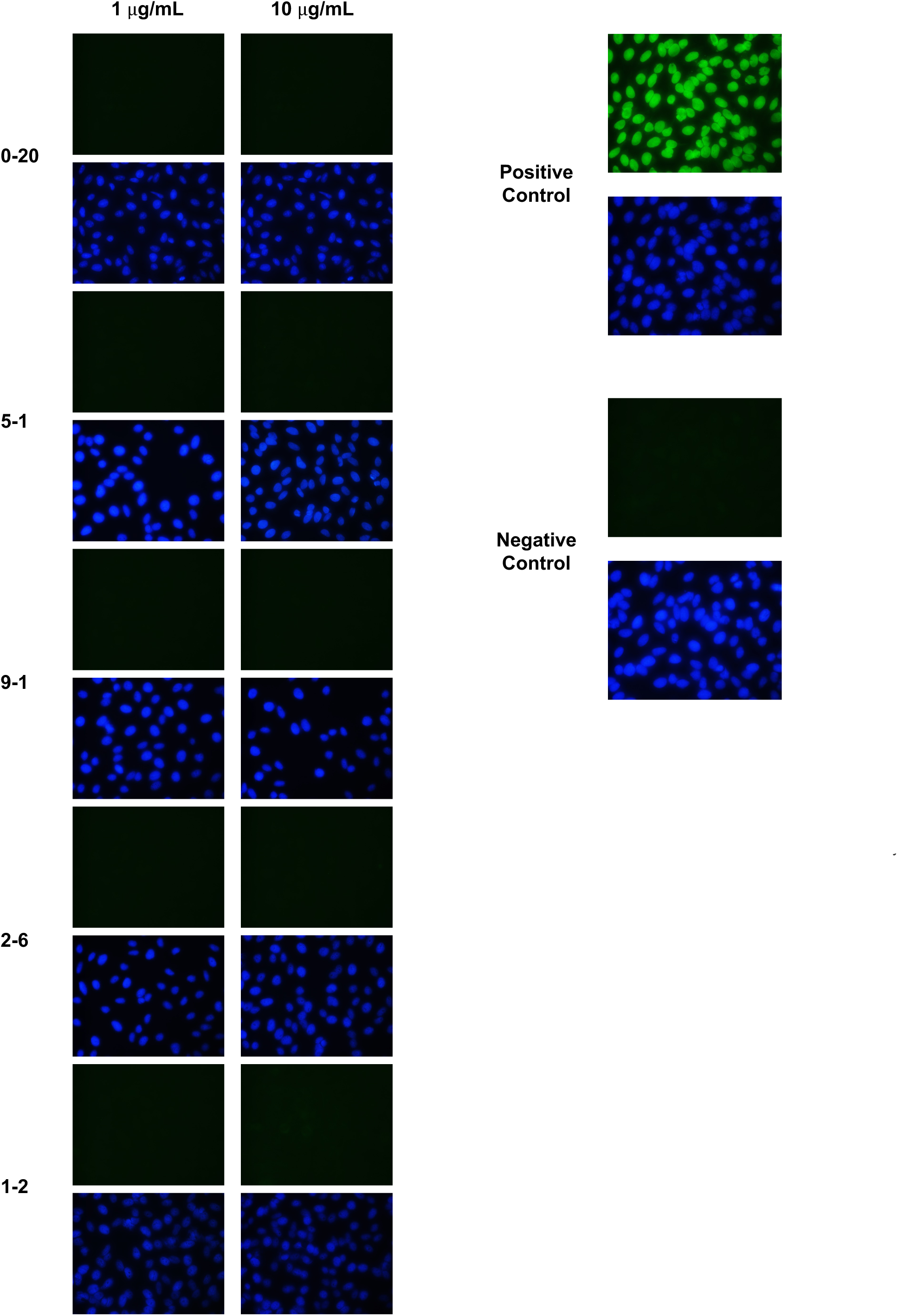
mAb binding to HEp2 Cells. Images of representative mAbs staining of HEp-2 cells. Indirect immunofluorescence assay testing reactivity of RSV/hMPV mAbs in HEp-2 cells. Each mAb was tested at 1 and 10 μg/mL. Positivity scores were determined relative to positive (ANA+ human serum) and negative (ANA – human serum) controls. DAPI staining (blue) was used to visualize nuclear DNA, goat anti-human Ig-FITC (green) staining notes Hep-2 cell reactivity. For all images, brightness was set to 150 and contrast was set to 100 using Photoshop.

**Supplementary Figure 2:**
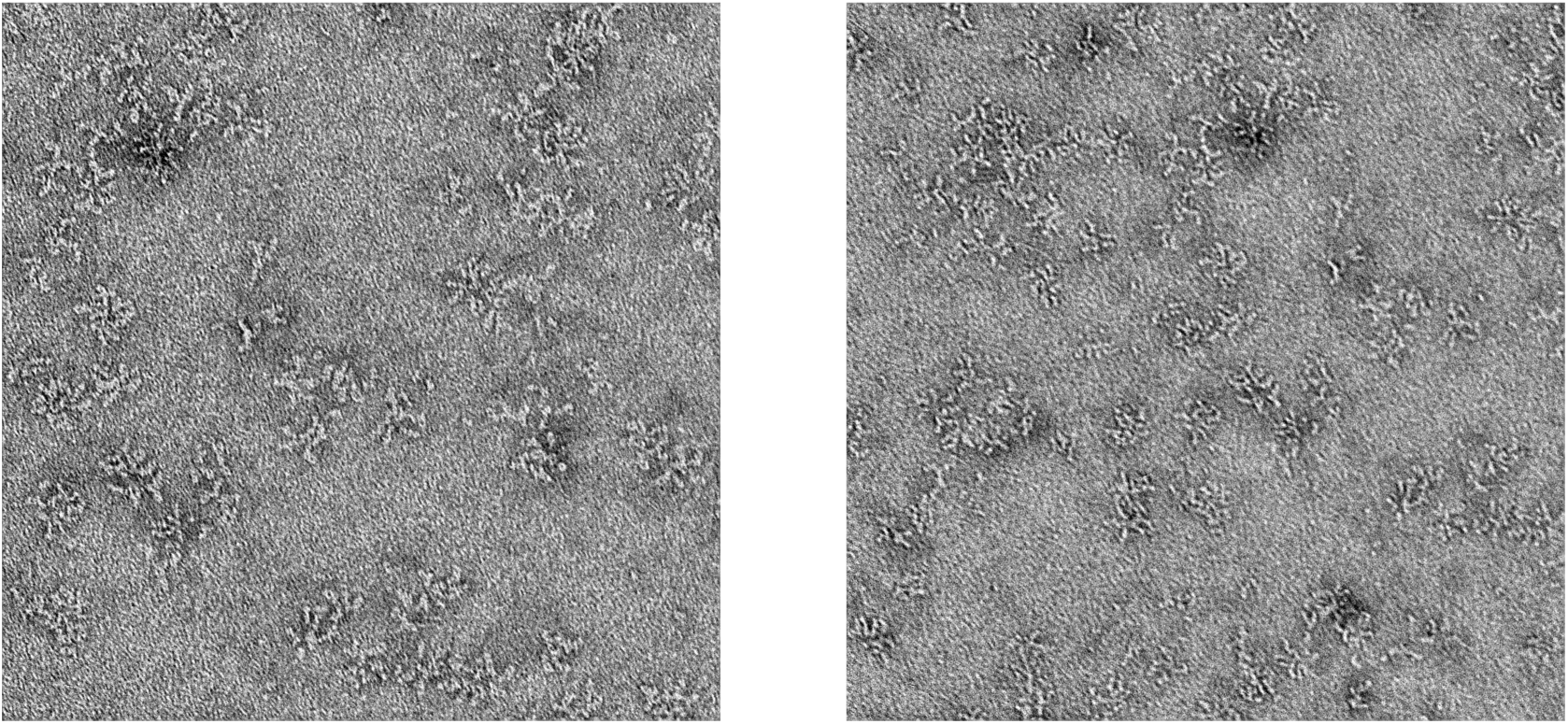
5-1 fab binding to DS-Cav1. DS-Cav1 complexed with 5-1 at 10 μg/ml. Left at 30 nm, right at 50 nm

**Supplementary Figure 3:**
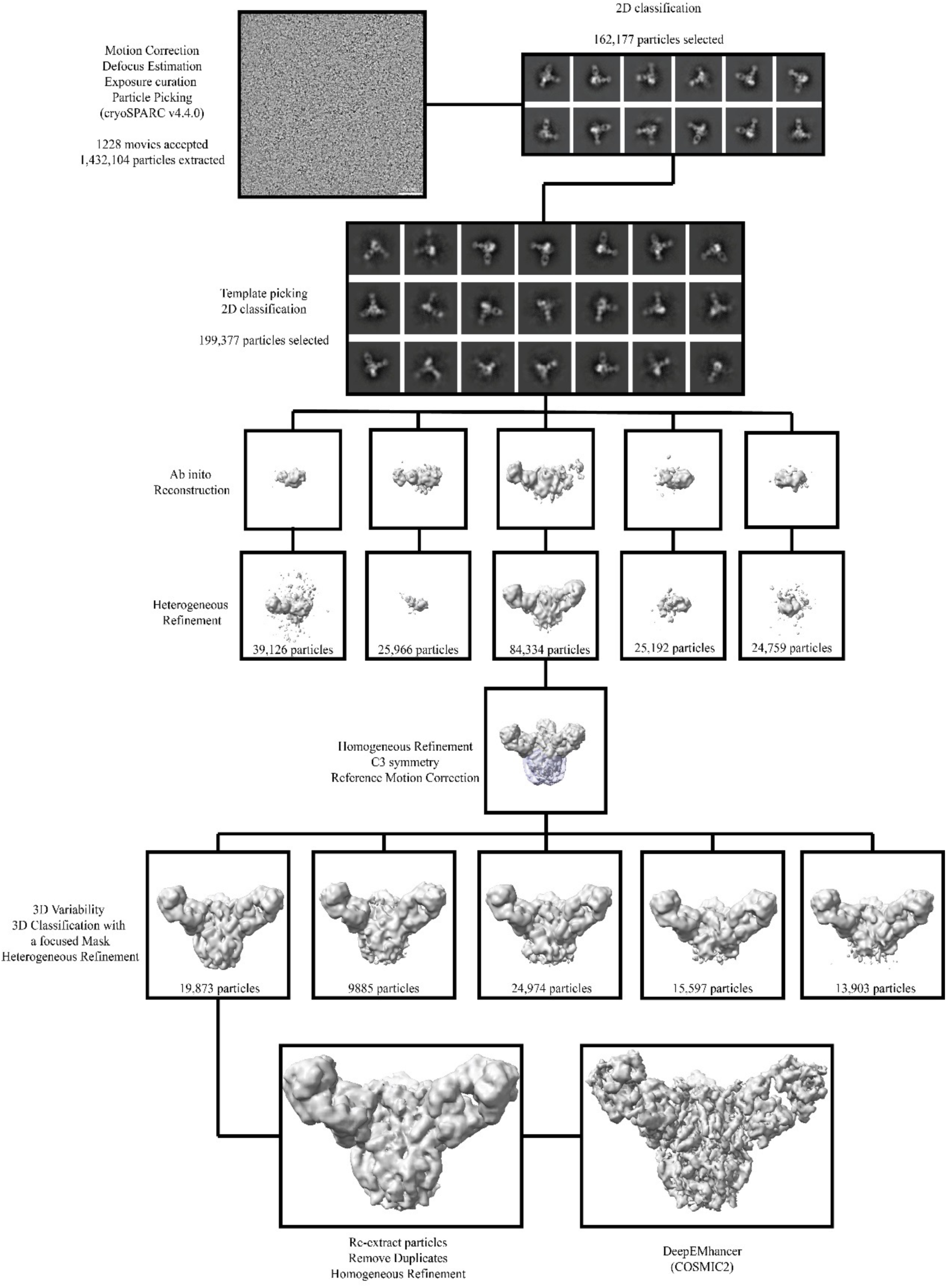
hMPV F and 5-1 Fab cryoEM dataset processing workflow. Representative micrographs, EM maps, computational programs and softwares from each step of the workflow are shown and labeled. The mask used for 3D classification is shown as a transparent purple surface.

**Supplementary Figure 4:**
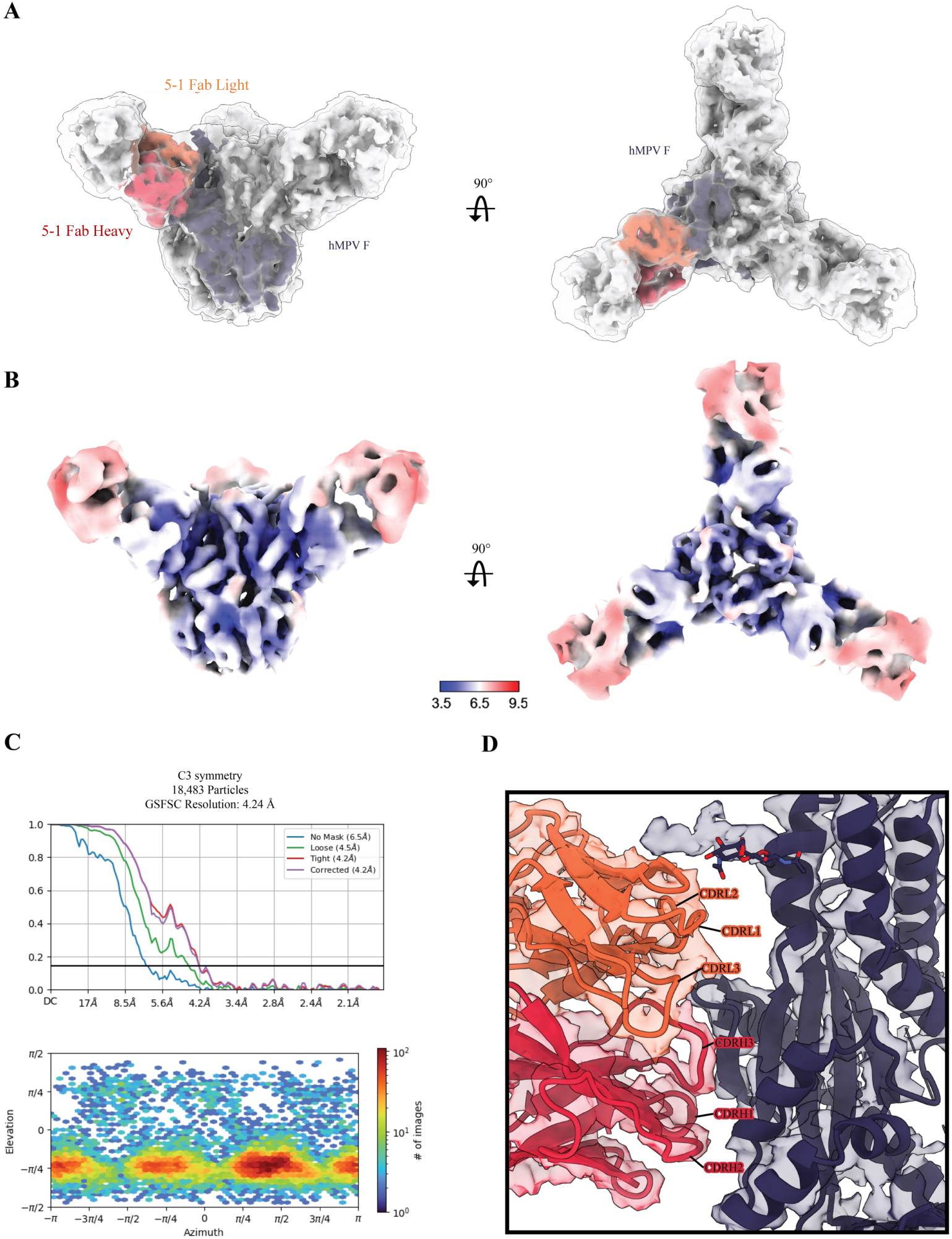
Validation of the obtained hMPV F EM map. A: Fitting of the DeepEMhanced EM map into the raw, unsharpened EM map. The raw, unsharpened EM map is shown as a transparent surface at the threshold of 0.0658. The DeepEMhanced EM map is shown as an opaque surface at the threshold of 0.431 with an individual hMPV F and Fab variable domain colored as indicated. B: The surface of the raw unsharpened EM map was colored by local resolution at the threshold of 0.026. C: FSC curves and particle orientation distribution for the EM map from the final homogeneous refinement step. Top, FSC curves; Botton, particle orientation distribution. Horizon line in FSC curves corresponds to an FSC value of 0.143. D. The binding interface between hMPV F-DsCavEs2-IPDS and 5-1 Fab. CryoEM map was shown as a transparent surface with the model fitted and colored.

**Supplementary Figure 5:**
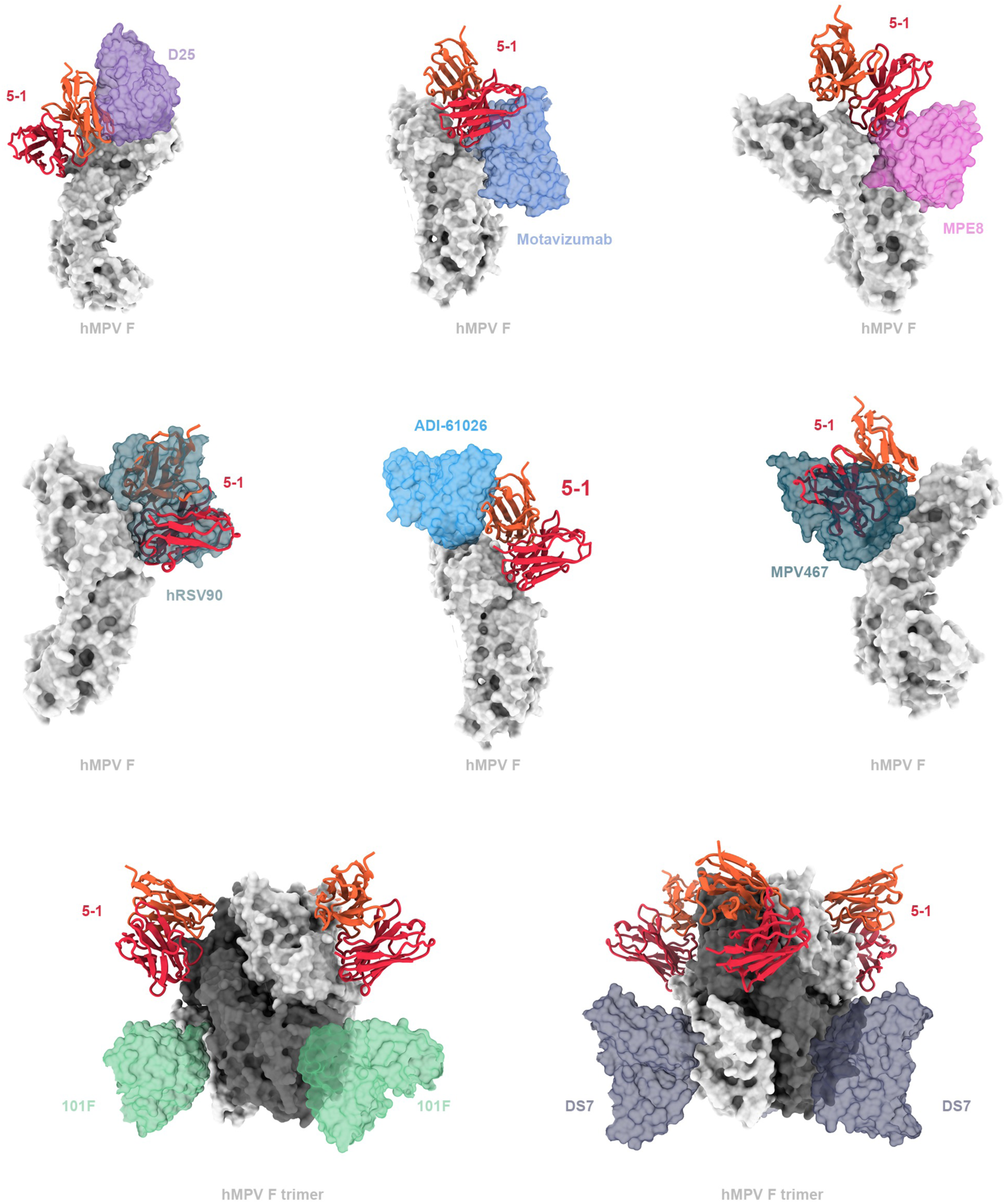
Steric clashes between 5-1 and site-specific antibodies. 5-1 shows significant clashes with competing antibodies and little to no steric clashing with non-competing antibodies from figure 2. Selected antibodies are shown as transparent surface and 5-1 is shown as cartoon with the light and heavy chain colored as orange and red, respectively.101 F and DS7 are modeled onto hMPV F trimers because of their close distance on native protomers.

**Supplementary Figure 6:**
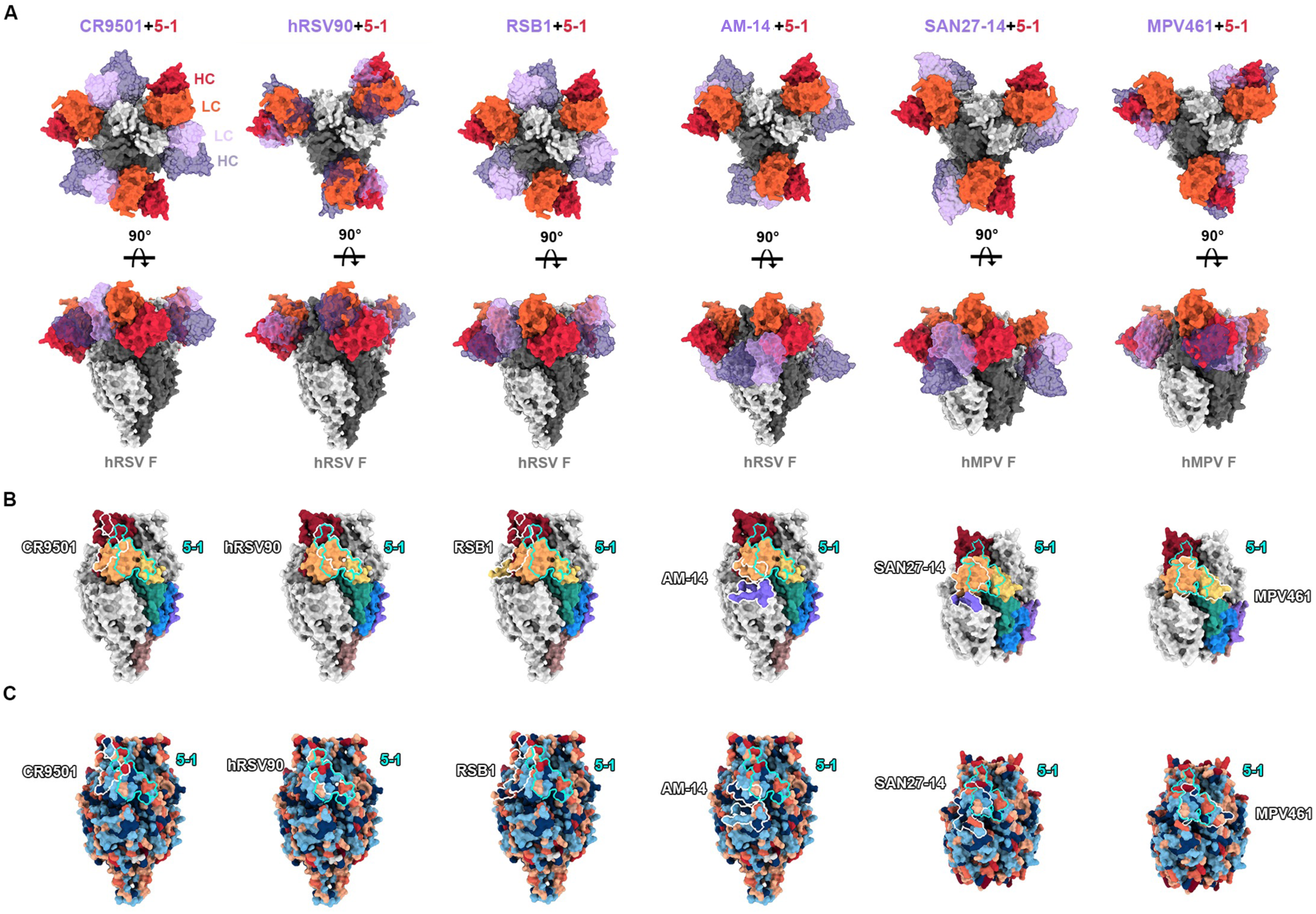
Binding poses and epitope conservation of antibodies binding site. **V** A: Modelling of site V antibodies with 5-1 Fab shows different binding poses on hMPV F. The quaternary antibody AM-14 was included for completeness. 5-1 is shown as opaque surface with the light and heavy chains colored as orange and red, respectively. Selected antibodies are modelled as transparent surface with the light and heavy chains colored as lavender and purple, respectively. B. Antigenic footprints of 5-1 and site V antibodies target different epitopes inside site V and often bind residues beyond site V. C. Comparison of epitopes based on sequence conservation reveals that sequence conservation did not solely determine the cross-neutralization properties of antibodies.

**Supplementary Figure 7:**
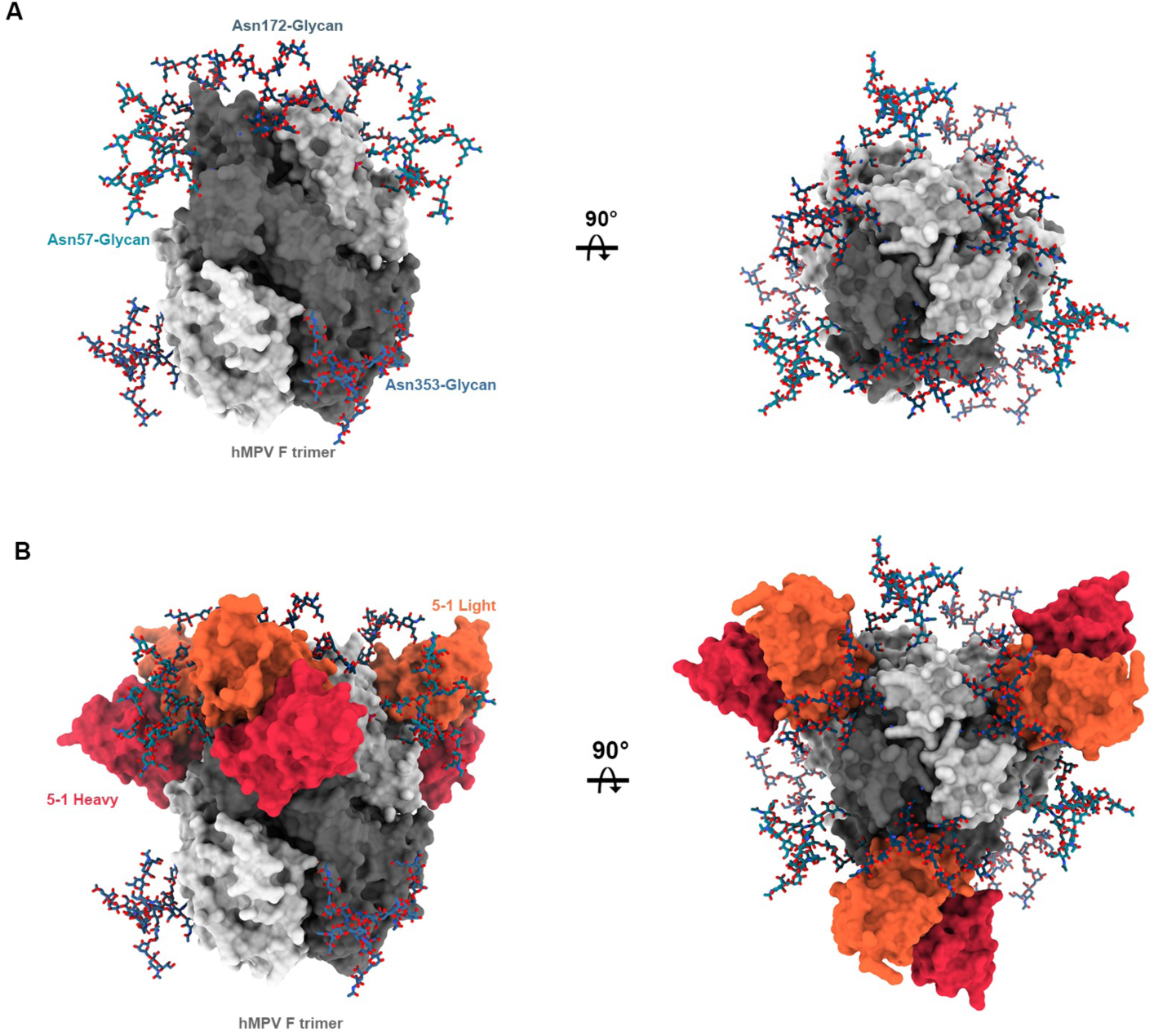
N-linked glycans and 5-1 Fab binding. A: Front view (left) and top view (right) of the N-linked complex glycans on hMPV F trimers. Glycans shown as ticks. B. Fit of the 5-1 Fab onto the modeled hMPV F trimers shows the light chain of 5-1 inserts into the cleft between Asn57-glycan and Asn172-glycan without clashes with Asn172-Glycan.

**Supplementary Table 1:**
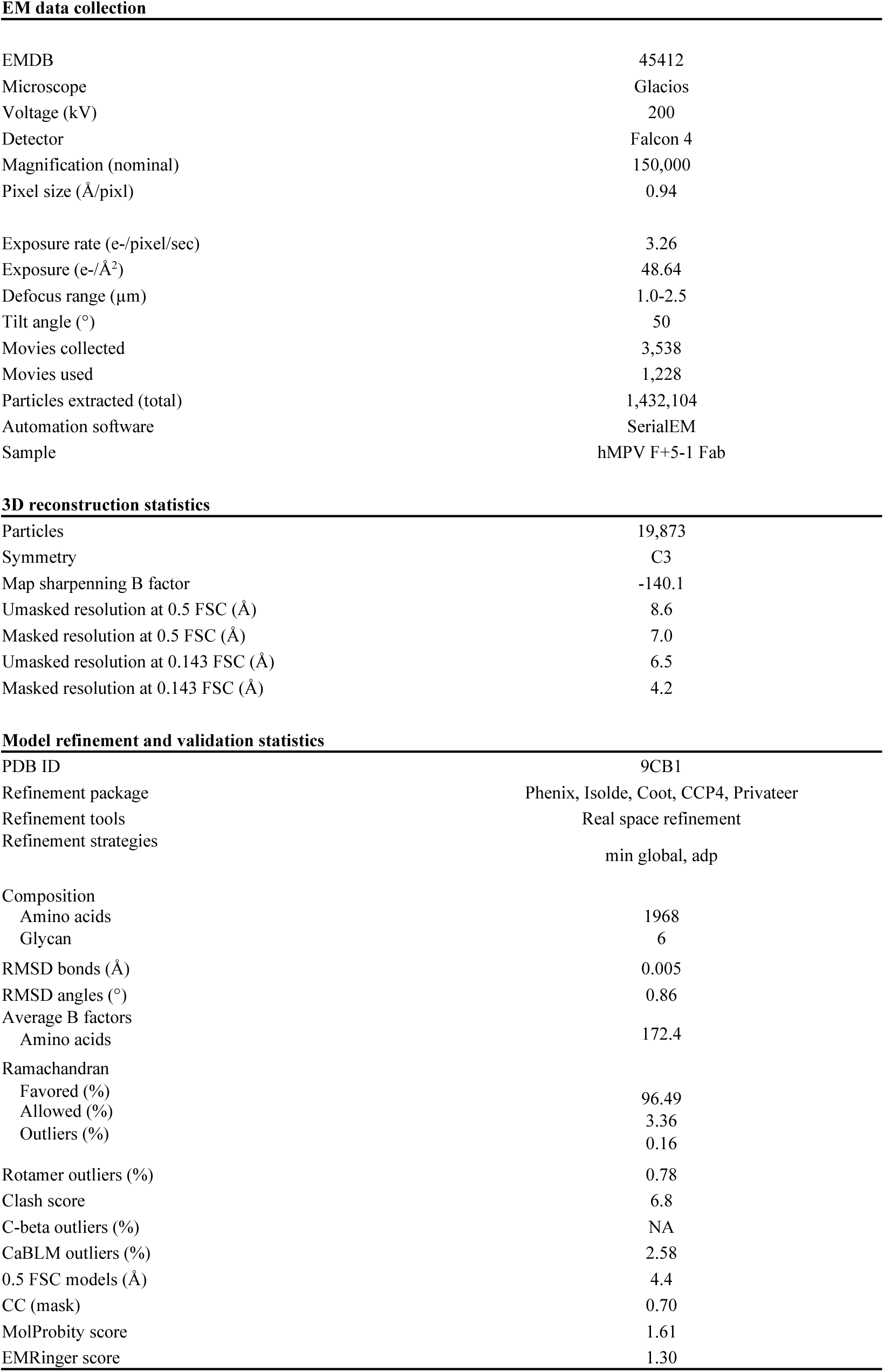
Cryo-EM data collection and reconstruction statistics.

**EM data collection**
|  |  |
| --- | --- |
| EMDB | 45412 |
| Microscope | Glacios |
| Voltage (kV) | 200 |
| Detector | Falcon 4 |
| Magnification (nominal) | 150,000 |
| Pixel size (Å/pixl) | 0.94 |
| Exposure rate (e-/pixel/sec) | 3.26 |
| Exposure (e-/Å<sup>2</sup>) | 48.64 |
| Defocus range (µm) | 1.0-2.5 |
| Tilt angle (°) | 50 |
| Movies collected | 3,538 |
| Movies used | 1,228 |
| Particles extracted (total) | 1,432,104 |
| Automation software | SerialEM |
| Sample | hMPV F+5-1 Fab |
| **3D reconstruction statistics** |  |
| Particles | 19,873 |
| Symmetry | C3 |
| Map sharpening B factor | -140.1 |
| Unmasked resolution at 0.5 FSC (Å) | 8.6 |
| Masked resolution at 0.5 FSC (Å) | 7.0 |
| Unmasked resolution at 0.143 FSC (Å) | 6.5 |
| Masked resolution at 0.143 FSC (Å) | 4.2 |

|  |  |
| --- | --- |
| PDB ID | 9CB1 |
| Refinement package | Phenix, Isolde, Coot, CCP4, Privateer |
| Refinement tools | Real space refinement |
| Refinement strategies | min global, adp |
| Composition |  |
| Amino acids | 1968 |
| Glycan | 6 |
| RMSD bonds (Å) | 0.005 |
| RMSD angles (°) | 0.86 |
| Average B factors |  |
| Amino acids | 172.4 |
| Ramachandran |  |
| Favored (%) | 96.49 |
| Allowed (%) | 3.36 |
| Outliers (%) | 0.16 |
| Rotamer outliers (%) | 0.78 |
| Clash score | 6.8 |
| C-beta outliers (%) | NA |
| CaBLM outliers (%) | 2.58 |
| 0.5 FSC models (Å) | 4.4 |
| CC (mask) | 0.70 |
| MolProbity score | 1.61 |
| EMRinger score | 1.30 |

